# Effects of LSD on music-evoked brain activity

**DOI:** 10.1101/153031

**Authors:** Mendel Kaelen, Romy Lorenz, Frederick Barrett, Leor Roseman, Csaba Orban, Andre Santos-Ribeiro, Matthew B Wall, Amanda Feilding, David Nutt, Suresh Muthukumaraswamy, Robin Carhart-Harris, Robert Leech

## Abstract

Music is a highly dynamic stimulus, and consists of distinct acoustic features, such as pitch, rhythm and timbre. Neuroimaging studies highlight a hierarchy of brain networks involved in music perception. Psychedelic drugs such as lysergic acid diethylamide (LSD) temporary disintegrate the normal hierarchy of brain functioning, and produce profound subjective effects, including enhanced music-evoked emotion. The primary objective of this study was to investigate the acute effects of LSD on music-evoked brain-activity under naturalistic music listening conditions. 16 healthy participants were enrolled in magnetic resonance imaging (fMRI) while listening to a 7-minute music piece under eyes-closed conditions on two separate visits (LSD (75 mcg) and placebo). Dynamic time courses for acoustic features were extracted from the music excerpts, and were entered into subject-level fMRI analyses as regressors of interest. Differences between conditions were assessed at group level subsequently, and were related to changes in music-evoked emotions via correlation analyses. Psycho-physiological interactions (PPIs) were carried out to further interrogate underlying music-specific changes in functional connectivity under LSD. Results showed pronounced cortical and subcortical changes in music-evoked brain activity under LSD. Most notable changes in brain activity and connectivity were associated with the component timbral complexity, representing the complexity of the music’s spectral distribution, and these occurred in brain networks previously identified for music-perception and music-evoked emotion, and showed an association with enhanced music-evoked feelings of wonder under LSD. The findings shed light on how the brain processes music under LSD, and provide a neurobiological basis for the usefulness of music in psychedelic therapy.

## 1. Introduction

The capacity of music to elicit emotions has been utilised by humans across cultures for millennia (Morley, 2013; Nettl, Bruno, 1956). Music is a highly dynamic and multi-dimensional stimulus, consisting of distinct acoustic features, such as pitch, loudness, tempo, rhythm and timbre. Within the field of cognitive neuroscience, the study of music has provided valuable insights into the functional organization of the human brain due to music’s recruitment of large-scale brain networks involved in attention (Janata, 2005), working memory (Kumar et al., 2016; Zatorre et al., 1994), episodic memory (Janata, 2009; Watanabe et al., 2008), motor function (Zatorre et al., 2007) and emotion (Koelsch, 2014; Trost et al., 2012). As such, neuroimaging studies with music highlight a hierarchy of processes involved in music perception: ascending from the cochlear, the auditory system analyses increasingly complex acoustic features which are subsequently integrated via multi-modal and higher-order cortical regions that facilitate associations with mental representations, memories and emotions (Kumar et al., 2007; Stewart et al., 2008). While the majority of neuroimaging studies have used controlled auditory experiments to study acoustic features in isolation, recent studies advocate the use of more naturalistic paradigms in which participants listen to real, extended pieces of music to increase the ecological validity and obtain a more comprehensive description of the brain mechanisms involved in processing naturalistic stimuli (Alluri et al., 2012; Burunat et al., 2016).

Advances in understanding the human brain can be made not only by studying its normal functioning, but also by experimentally perturbing it. For the study of music, perturbations that may be of particular significance are the effects of classic psychedelic drugs such as lysergic acid diethylamide (LSD) and psilocybin (i.e. psychedelics). Psychedelics activate the serotonin 2A receptor (Glennon et al., 1984; Titeler et al., 1988; Vollenweider et al., 1998), and produce an altered state of consciousness that is characterised by marked reductions in functional coupling within high-level brain networks (Carhart-Harris et al., 2012, 2016a; Palhano-Fontes et al., 2015), but simultaneous increased cross-talk between low-level areas (Tagliazucchi et al., 2014, 2016). This network “collapse” is thought to be a core mechanism underlying their subjective effects, that include a dissolving of the usual sense of self-hood, increased emotionality and vivid mental imagery (Carhart-Harris et al., 2014; Lebedev et al., 2015) and a profoundly altered subjective experience of music (Kaelen et al., 2015).

Studying psychedelics and music in combination may therefore represent a unique opportunity to improve our understanding of changes in the brain’s processing of complex naturalistic stimuli, accompanying the changes in functional brain hierarchies that are reported under psychedelics, and to investigate how these changes are related to reported changes in the subjective experience of music. In addition, increasing numbers of studies assess the therapeutic potential of these subjective effects within “psychedelic therapy” - a therapeutic approach that is characterised by lengthy periods of introspective music-listening during drug effects (Carhart-Harris et al., 2016b; Gasser et al., 2014; Griffiths et al., 2016; Grob CS et al., 2011; Johnson et al., 2014; Ross et al., 2016). Despite the prominence of music in psychedelic therapy, the effects of psychedelics on the subjective experience of music have only been empirically investigated recently (Kaelen et al., 2015, 2016). Studying how psychedelics and music work together in the brain, may therefore also be important to advance our understanding of the use of music in psychedelic therapy.

The primary objective of this study was to investigate the acute effects of the classic psychedelic LSD on music-evoked brain activity under naturalistic music listening conditions. In addition, music can reliably evoke emotions (Trost et al., 2012), and psychedelic drugs are shown to reliably facilitate emotionally intense “peak-experiences”^1^ under music-listening conditions (Griffiths et al., 2011, 2006). Hence, the secondary objective of the study was to investigate the neural correlates of “peak” emotions such as feelings of *wonder* and *transcendence* evoked by music under LSD. These objectives were studied as part of a larger project examining the effects of LSD on brain activity in healthy volunteers using multimodal neuroimaging (reported elsewhere (Carhart-Harris et al., 2016a; Lebedev et al., 2016; Tagliazucchi et al., 2016)).

## 2. Materials and Methods

### 2.1 Ethical approvals

The study received approvals by the National Research Ethics Service (NRES) committee London – West London and was conducted in accordance with the revised declaration of Helsinki (2000), the International Committee on Harmonisation Good Clinical Practice guidelines and National Health Service (NHS) Research Governance Framework. Imperial College London sponsored the study, which was conducted under a Home Office license for research with schedule I drugs.

### 2.2 Participants

Twenty participants (16 males and 4 females) were recruited for the study after successful screening and written informed consent. Screening consisted of physical and mental health assessments. Physical health assessments included electrocardiogram (ECG), routine blood tests, disclosure of full medical history and urine test for recent drug use and pregnancy. Mental health assessment consisted of psychiatrics interview. All participants provided full disclosure of their drug use history. Exclusion criteria included: being younger than 21 years of age, having a personal history of diagnosed psychiatric illness, having an immediate family history of a psychotic disorder, having used illicit drugs within 6 weeks of the first scanning day, having experienced a persistent adverse reaction to a psychedelic drug, pregnancy, problematic alcohol-use (i.e. > 40 units consumed per week), and having a medical condition rendering them unsuitable for the study. All participants were required to have at least one previous experience with a classic psychedelic drug (e.g. LSD, mescaline, psilocybin or dimethyltryptamine (DMT)). One participant did not complete the MRI scans due to anxiety and three participants were excluded from analyses due to technical problems with the sound delivery. This resulted in the data of 16 participants (13 males, 3 female) being included in the data-analysis; with preserved counterbalanced experimental design.

### 2.3 Stimuli

Two excerpts (A and B) from two songs by ambient artist Robert Rich were selected for the study. The stimuli were both 7.3 min long and were balanced in their acoustic properties, and rich in timbre, but not in rhythm (see Appendix: Table 1 and Figure 1). Pre-study assessments in a separate group confirmed balance for their emotional potency. To reduce interference of fMRI scanning noise with the music experience, volume-maximization and broadband compression was carried out using Ableton live 9 software. Each participant listened to both stimuli, in a balanced order across conditions.

**Figure 1.**
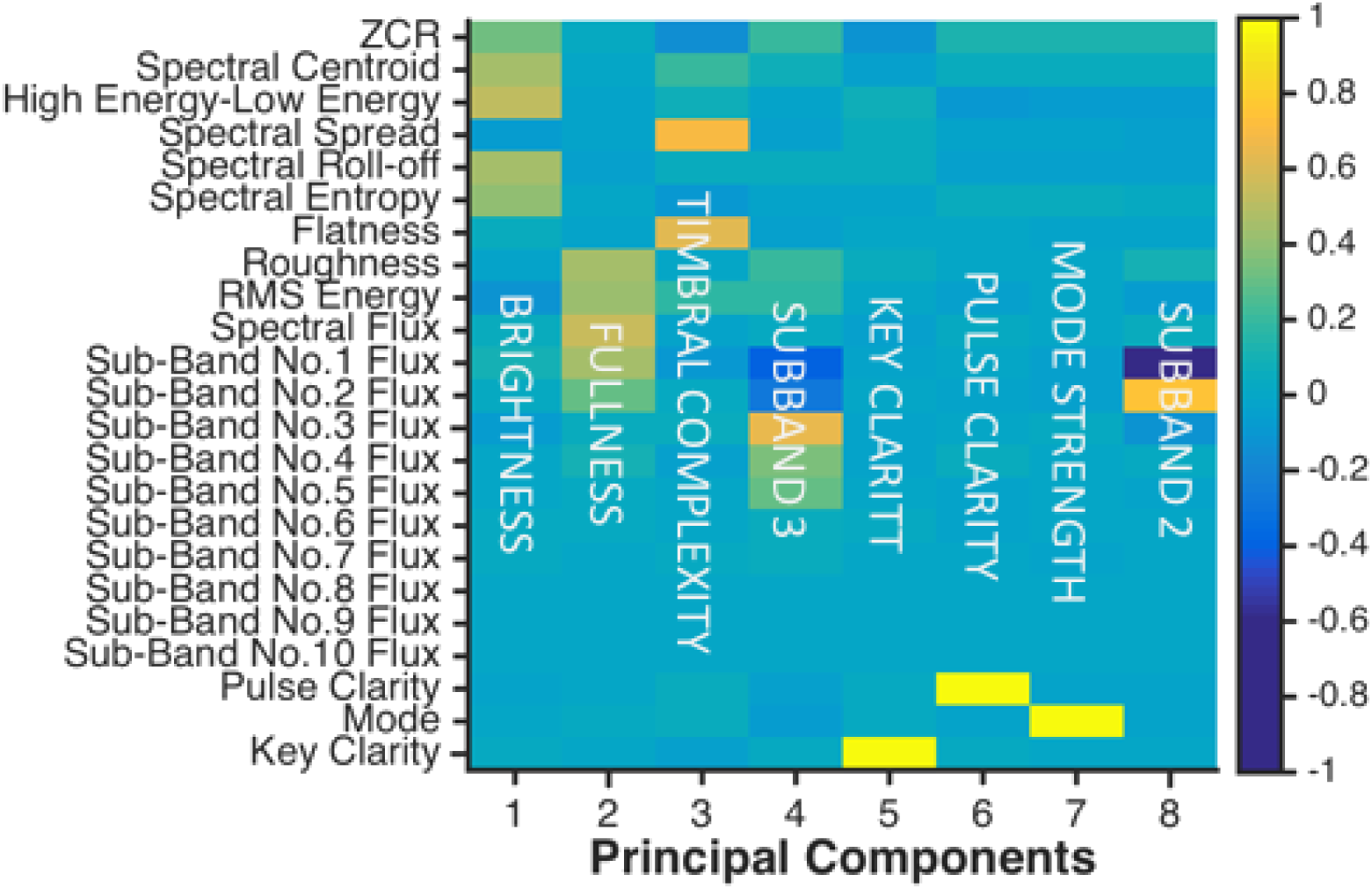
Principle Component Analysis (PCA) of acoustic features. Loadings of the acoustic features on the first 8 PCs obtained from PCA followed by varimax rotation explained more than 90% of the variance. The x-axis shows the ordering of principal components, with the components ordered by explained variance. The colour bar corresponds to the strength of the loading for each acoustic feature for that components: warm colours indicates a positive loading, and cold colours a negative loading.

**Table 1.** Acoustic features from the Music Information Retrieval toolbox that were used in the analysis.

|  |
| --- |
| **Short-term features** |
| Zero crossing rate, spectral centroid, high energy–low energy ratio, spectral spread, spectral roll-off, spectral entropy, spectral flatness (Wiener entropy), roughness, RMS energy, spectral flux, and Sub-Band Flux (10 coefficients). |
| **Long-term features** |
| Pulse clarity, fluctuation centroid, fluctuation entropy, mode and key clarity. |

### 2.4 Experiment overview and procedures

All participants attended two study days that were separated by at least 14 days. LSD was received on one of the study days, and placebo (saline) on the other. The condition-order was balanced across participants, and participants, but not the researchers, were blind to the condition-order.

All participants received 75 μg of LSD intravenously via a 10ml solution infused over a two-minute period via an inserted cannula. Administration was always followed by an acclimatization period of approximately 60 min, in which participants were encouraged to relax. Drug effects were noticed between 5 to 15 min post-dosing, approached maximum intensity between 60 to 90 min post-dosing, and maintained peak intensity for approximately four hours post-dosing. Blood-Oxygen-Level Dependent (BOLD) MRI scanning was performed during peak drug intensity, starting approximately 120 minutes post-dosing, and lasted for approximately 60 minutes. Once the subjective effects of LSD had sufficiently subsided, the study psychiatrist assessed the participant’s suitability for discharge.

Each fMRI scanning session involved three eyes-closed resting state scans, each lasting seven minutes and twenty seconds. The music-listening scan (220 TRs, 7.3 min) always occurred after the first resting state (no music) scan and before a final resting-state scan (no music). Prior to each scan, participants were instructed via a display screen to close their eyes and relax. The music itself was triggered by the first TR, and listened to via MR compatible headphones (MR Confon). Music volume was adjusted to a personal level that was “loud, but not unpleasant” and maintained for both conditions.

### 2.5 Qualitative measurements

The intensity of different types of music-evoked emotion was assessed via the 25-item Geneva emotional music scale (GEMS)(Zentner et al., 2008), and was administered under placebo and under LSD immediately after fMRI scanning was completed. The GEMS assesses music-evoked or directly “personally felt” emotion, and not the perception of musical emotion in the music. For example: a piece of music may sound sad to the listener, but the listener may not necessarily feel sad him or herself in response to this music. The GEMS is designed to assess the latter. The 25 items of the GEMS were averaged into nine distinct factors of emotions: wonder, transcendence, power, tenderness, nostalgia, peacefulness, joyful activation, sadness, and tension. A paired t-test was performed to compare ratings between conditions (LSD versus placebo) for each emotion. Non-parametric permutation testing (10,000 permutations, two-tailed) was carried out to correct for multiple comparisons.

### 2.6 Acoustic feature extraction

Acoustic features from excerpt A and B were extracted separately using the Music Information Retrieval (MIR) toolbox (version 1.6.1), implemented in MATLAB (Lartillot and Toiviainen, 2007). Selection of acoustic features was based on Alluri et al (2012, 2013), who identified 25 features that broadly capture the dynamics of the timbral, tonal and rhythmic properties present in the music stimuli (Alluri et al., 2012, 2013). Following Alluri et al, from the original 25 features, we excluded fluctuation entropy and fluctuation centroid because they have not been studied in the literature and did not carry any valid perceptual meaning (Alluri et al. 2012, Alluri 2016 personal communication). The 23 acoustic features used in our analyses were categorized into short-term features and long-term features (for an overview see Table 1). The short-term features were extracted with a sliding window of 25 ms, with 50% overlap between each segment while for the computation of the long-term features a longer sliding window of 3 s was employed, with 33% overlap between each segment (following the approach in Alluri et al. 2012, 2013). Subsequently, principle component analysis (PCA) on the acoustic features (combined for excerpt A and B) was performed to (1) identify acoustic properties that are present in both excerpts and (2) to reduce the number of regressors entered into the general linear model (GLM) for fMRI analyses. To prepare the acoustic features for the PCA, we first resampled short and long-term features to the same sampling rate of 1 s. We then rescaled (between 0 and 1) the acoustic features. The resulting timecourses for excerpt A and B were then concatenated, to undergo the PCA. We performed varimax rotation on the first eight principal components (PCs) that explained over 90% of the variance (see Fig. 1 for rotated PCs and their respective loadings).

In order to validate that the first eight PCs explained the two excerpts equally well (i.e., that PCs were not driven by either one excerpt or the other), we correlated (Pearson r) each of the 23 predicted timeseries (by multiplying the eight rotated PC loadings with the 23 PC timecourses) with the corresponding acoustic features for the two excerpts separately. Across all 23 acoustic features, we found a mean ± std r of 0.9098 ± 0.0919 for excerpt A and 0.8835 ± 0.1037 for excerpt B (for correlation values per acoustic feature, see Appendix Fig. 1). As the difference in variance explained between the two excerpts was negligible, we used the timecourses of the first eight rotated components for our subsequent analyses.

Next, we split the timecourses of the rotated PCs to obtain timeseries corresponding to excerpt A and B, respectively. Those were then convolved with a double-gamma hemodynamic response function (HRF) to account for the hemodynamic delay, down-sampled to 2 s, to match the sampling rate of the fMRI data and high-pass filtered at the cut-off frequency of 0.01 Hz (same as fMRI data). The first 10 seconds (5 TRs) of each excerpt’s timeseries were excluded for subsequent analysis in order to avoid any artefacts caused by the convolution operation. The timeseries were then entered as regressors of interest into subject-level GLMs of the fMRI analysis.

### 2.7 MRI scanning

Neuroimaging was performed on a 3T GE HDx system. An initial 3D FSPGR anatomical scan was obtained in an axial orientation, with field of view = 256× 256× 192 and matrix = 256 × 256× 192 to yield 1-mm isotropic voxel resolution (TR/TE = 7.9/3.0 ms; inversion time = 450 ms; flip angle = 20°). This was used for registration and segmentation of functional images. Functional images were acquired using a gradient echo planar imaging sequence, TR/TE = 2000/35ms, field-of-view = 220mm, 64 × 64 acquisition matrix, parallel acceleration factor = 2, 90° flip angle. 35 oblique axial slices were acquired in an interleaved fashion, each 3.4mm thick with zero slice gap (3.4mm isotropic voxels). The precise length of each of the BOLD scans was 7 minutes and 20 seconds.

### 2.8 fMRI pre-processing

The fMRI pre-processing pipeline was assembled from four complementary imaging software packages. Specifically, FMRIB Software Library (FSL) (Smith et al., 2004), AFNI (Cox, 1996), Freesurfer (Dale et al., 1999) and Advanced Normalization Tools (ANTS) (Avants et al., 2009) were used. The following pre-processing stages were performed: 1) de-spiking (3dDespike, AFNI); 2) motion correction (3dvolreg, AFNI) by registering each volume to the volume most similar, in the least squares sense, to all others (in-house code); 3) brain extraction (BET, FSL); 4) rigid body registration to anatomical scans (eleven subjects with FSL’s BBR, one subject with Freesurfer’s bbregister and four subjects manually); 5) non-linear registration to 2mm MNI brain (Symmetric Normalization (SyN), ANTS); 6) scrubbing (Power et al., 2012) - using a framewise displacement (FD) threshold of 1.0 mm; 7) spatial smoothing (FWHM) of 5mm (3dBlurInMask, AFNI) and high-pass filtering at a cut-off frequency of 0.01 Hz to remove baseline signal drifts. In addition, nine nuisance regressors were obtained: six were motion-related (3 translations, 3 rotations) and three were anatomically-related (not smoothed). Specifically, the anatomical nuisance regressors were: I) ventricles (Freesurfer, eroded in 2mm space), II) draining veins (DV) (FSL’s CSF minus Freesurfer’s Ventricles, eroded in 1mm space) and III) white matter (WM) (FSL’s WM minus Freesurfer’s subcortical grey matter (GM) structures, eroded in 2mm space). In order to cut the fMRI data and nuisance regressors to the same length as the PC timecourses, we removed the first 5 TRs.

### 2.9 fMRI data analysis

We applied a standard analysis pipeline from FSL FEAT GLM to determine the effect of PC timecourses on BOLD activation. All eight PC timecourses were entered as regressors of interest into subject-level GLMs in addition to their temporal derivatives and all nine nuisance regressors (resulting in 27 regressors in total). We then analysed the fMRI data with respect to eight contrasts (one contrast for each PC). The resulting individual subject-level contrasts were then entered into a group averaged higher-level (mixed-effects) cluster-corrected FEAT analysis (FLAME 1+2) to obtain the paired t-test contrasts of LSD > Placebo and Placebo < LSD. In order to model out any potential effect of the order in which subjects listened to the different music excerpts under the different conditions, respectively, we included an interaction term in the higher-level FEAT as a regressor of no interest. We repeated the higher-level analysis without the interaction term and obtained qualitatively similar results (not reported). All final group-level images were thresholded using a cluster correction threshold of Z > 2.3 and a cluster significance threshold of p = 0.05. Group-level images were visualized on an average surface brain using MRIcron(Rorden et al., 2007).

### 2.10 ROI selection and extraction for correlation with individual variability in reported music-evoked peak emotions

Region of interest (ROI) analysis was constrained to the group-level contrast LSD>Placebo for *timbral complexity* (i.e., PC3, see Results). ROI selection was based on previous literature and encompassed structures that have been commonly identified for music-evoked emotion (striatum, precuneus and insula) (Koelsch, 2014; Trost et al., 2012), auditory perception (planum temporale and IFG) (Alluri et al., 2012, 2013; Griffiths and Warren, 2002; Zatorre and Salimpoor, 2013) and the subjective effects of psychedelics (precuneus and insula) (Carhart-Harris et al., 2016a; Muthukumaraswamy et al., 2013a; Tagliazucchi et al., 2016). ROIs were created by first selecting corresponding binarized structural areas from the Harvard-Oxford atlas (using a probability threshold of 10%) and then multiplying these with the group-level results to guarantee anatomical and functional specificity of the ROIs. Note that only the largest consecutive cluster of voxels was considered within each ROI. This procedure led to nine ROIs: right planum temporale, left planum temporale, right precunes, right putamen, left putamen, right caudate, left caudate, right inferior frontal gyrus and right anterior insula. The mean parameter estimates within each ROI were extracted for each participant, and Spearman’s rank correlation tests were used to relate the changes in parameter estimates (LSD > Placebo) with changes in GEMS-ratings for music-evoked emotions for the factors *wonder* and *transcendence* (LSD > Placebo). Non-parametric permutation testing (10,000 permutations, one-tailed) was carried out to correct for multiple comparisons (9 ROIs x 2 GEMS scales).

### 2.11 Psychophysiological interaction analysis

In order to assess *timbral*-specific changes in functional connectivity for the brain regions that showed a significant relationship with music-evoked feelings of wonder (i.e., precuneus and right inferior frontal gyrus, see Results), we conducted two separate psychophysiological interaction (PPI) analyses. For this, we first extracted subject-specific timecourses from those ROIs and then applied the standard FSL FEAT GLM analysis pipeline. In addition to regressors that were included in the initial fMRI analysis (eight PC timecourses, their temporal derivatives and nine nuisance regressors), we included the ROI timecourse (physiological regressor) and the PPI regressor that consisted of the element-wise multiplication of the physiological regressor with the timecourse encoding timbral complexity (psychological regressor, i.e. PC 3). The resulting individual subject-level contrasts were then entered into the same higher-level (mixed-effect) cluster-corrected FEAT analysis as described above (with an interaction term to model out the effect of “group”, see fMRI analysis section).

## 3. Results

### 3.1 Acoustic feature extraction

The first eight principal components that together explained more than 90% of the variance, showed consistency with the components identified in other musical genres using the same method (tango, classic and pop (Alluri et al., 2012, 2013; Burunat et al., 2016)), supporting their reliability for representing fundamental acoustic properties in music. Importantly, perceptual validation and labelling for similar components has been demonstrated (Alluri et al., 2012). In the present study, we therefore adopted a comparable labelling for the identified principle components (PCs) (Fig. 1). In fact, when quantitatively comparing our and Alluri et al.’s PCs, we find medium to very high correlations (Pearson r) for fullness (*r* = 0.37), brightness (*r* = .96), timbral complexity (*r* = .91), key clarity (*r* = .96) and pulse clarity (*r* = .96).

### 3.2 Effects of LSD on music-evoked brain activity

Following pre-processing (see Methods), the time courses of the first eight PCs were entered into subject-level fMRI analyses as regressors of interests. Resulting individual subject-level contrasts were then entered into high-level analyses to obtain paired t-test contrasts of LSD > Placebo and Placebo < LSD.

These analyses revealed altered BOLD responses under LSD for different acoustic components and within distinct cortical and sub-cortical areas. For the component *brightness*, decreases under LSD were observed in the left lateral occipital cortex (Fig. 2A). For *fullness,* LSD increased activation in the bilateral thalamus, and decreased activation in the occipital cortex and right middle and inferior frontal gyrus (IFG) (Fig. 2B). For *timbral complexity* widespread increases under LSD were found in cortical and subcortical networks, including auditory and auditory association cortices such as Heschl’s Gyrus, superior temporal gyrus (STG), and planum temporale as well as higher-level heteromodal cortices such as the precuneus, the right insula and the right IFG, bilateral striatum, as well as the occipital cortex and the supplementary motor cortex (see Supplementary Table 3 for a complete list). Increased BOLD activations to *timbral complexity* appeared more evident for left auditory cortices and more extensive in the right IFG and right anterior insula, although this was not directly compared (Fig. 2C). For the component *Subband 3,* LSD significantly increased activation in occipital cortices while decreasing activation in the cerebellum (Fig. 2D). For *key clarity*, significant increases under LSD were observed in the right lateral occipital cortex, and decreases were found in the left striatum (Fig. 2E). For the component *Subband 2,* LSD-related increases were observed in the right lateral occipital cortex (Fig. 2F).

**Figure 2.**
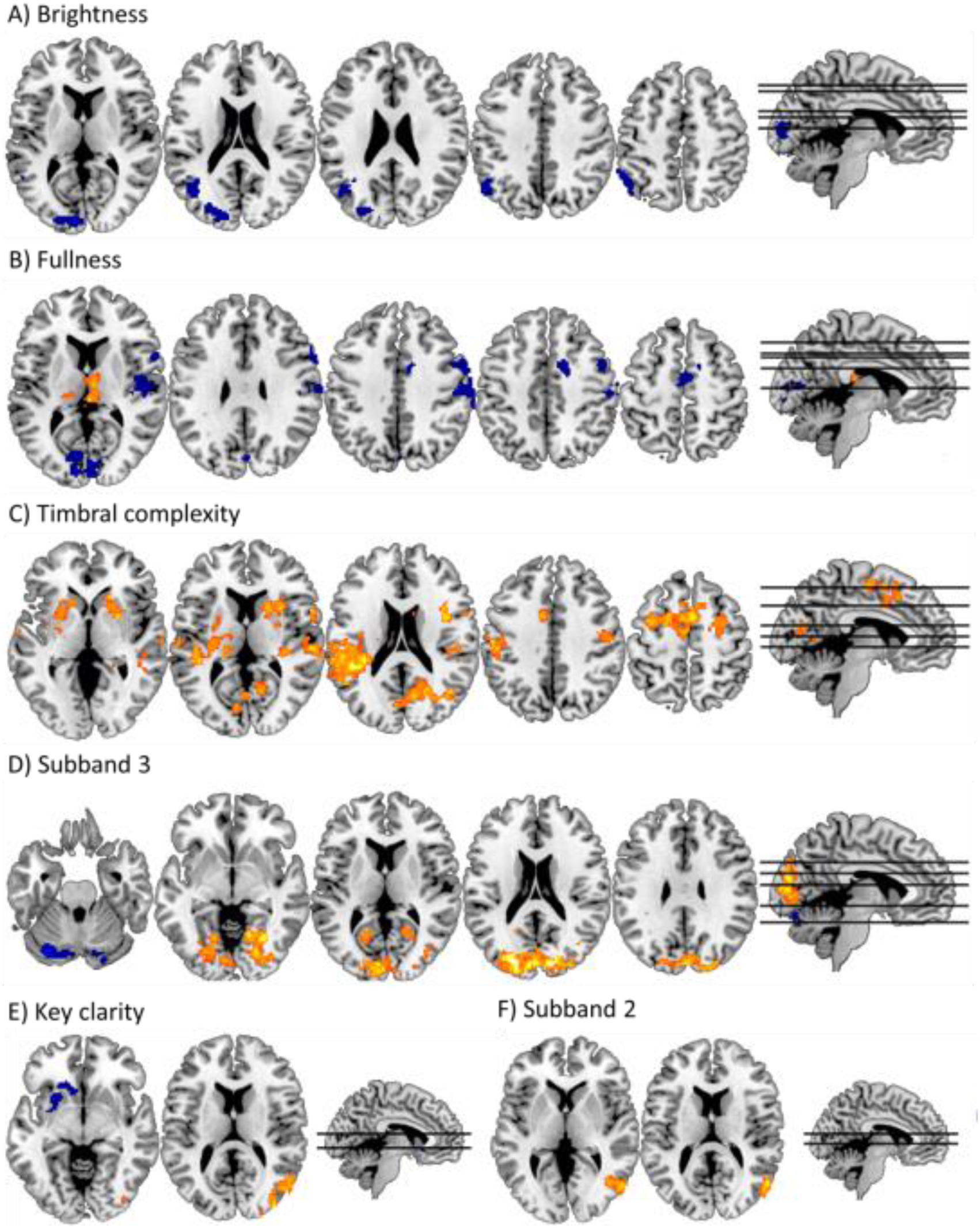
Brain regions showing altered BOLD response to the acoustic features under LSD. Increases (LSD>placebo) are displayed in yellow, and decreases (placebo>LSD) are displayed in blue. Cluster-correction was applied to all images with a threshold of Z > 2.3. Significant effects were observed in: **A)** *brightness*, with decreases in the occipital cortex, in **B)** *fullness*, with decreases in occipital cortex and right inferior frontal gyrus (IFG) and increases in bilateral thalamus, in **C)** *timbral complexity*, with increases in bilateral striatum, occipital cortex, supplementary motor cortex, auditory cortices, right insula and right IFG, in **D)** *subband 3*, with decreases in the cerebellum, and increases in the occipital cortice, in **E)** *key clarity*, with decreases in the left striatum and the right lateral occipital cortex, and finally, in **F)** *subband 2*, with increases in the right occipital cortex.

### 3.3 Effects of LSD on music-evoked emotion

Alongside investigating changes in music-evoked brain activity under LSD, our study also assessed the effects of LSD on music-evoked emotions. We measured nine different music-evoked emotions using the Geneva Emotional Music Scale (GEMS) that was completed by each participant after listening to music in each condition(Zentner et al., 2008). Paired t-tests on the GEMS scales revealed that, compared to placebo, music-evoked emotions were significantly higher under LSD for *wonder* (*t*_(18)_=4.47, *p*=.002), *transcendence* (*t*_(18)_=4.17, *p*=.004), *power* (*t*_(18)_=3.82, *p*=.008), *tenderness* (*t*_(18)_=3.78, *p*=.008), *nostalgia* (*t*_(18)_ = 3.94, *p* = .006), *peacefulness* (*t*_(18)_ = 4.45, *p*=.003), and *joyful activation* (*t*_(18)_=4.86, *p*=.001), but not for *sadness* (*t*_(18)_=1.93, *p*=.25) or *tension* (*t*_(18)_=1.42, *p*=.63) (Fig. 3). P-values reported here are corrected for multiple comparisons using two-sided permutation testing with α set to .05. These results replicate our previous work (Kaelen et al., 2015) demonstrating an intensified emotional experience with music under LSD.

**Figure 3.**
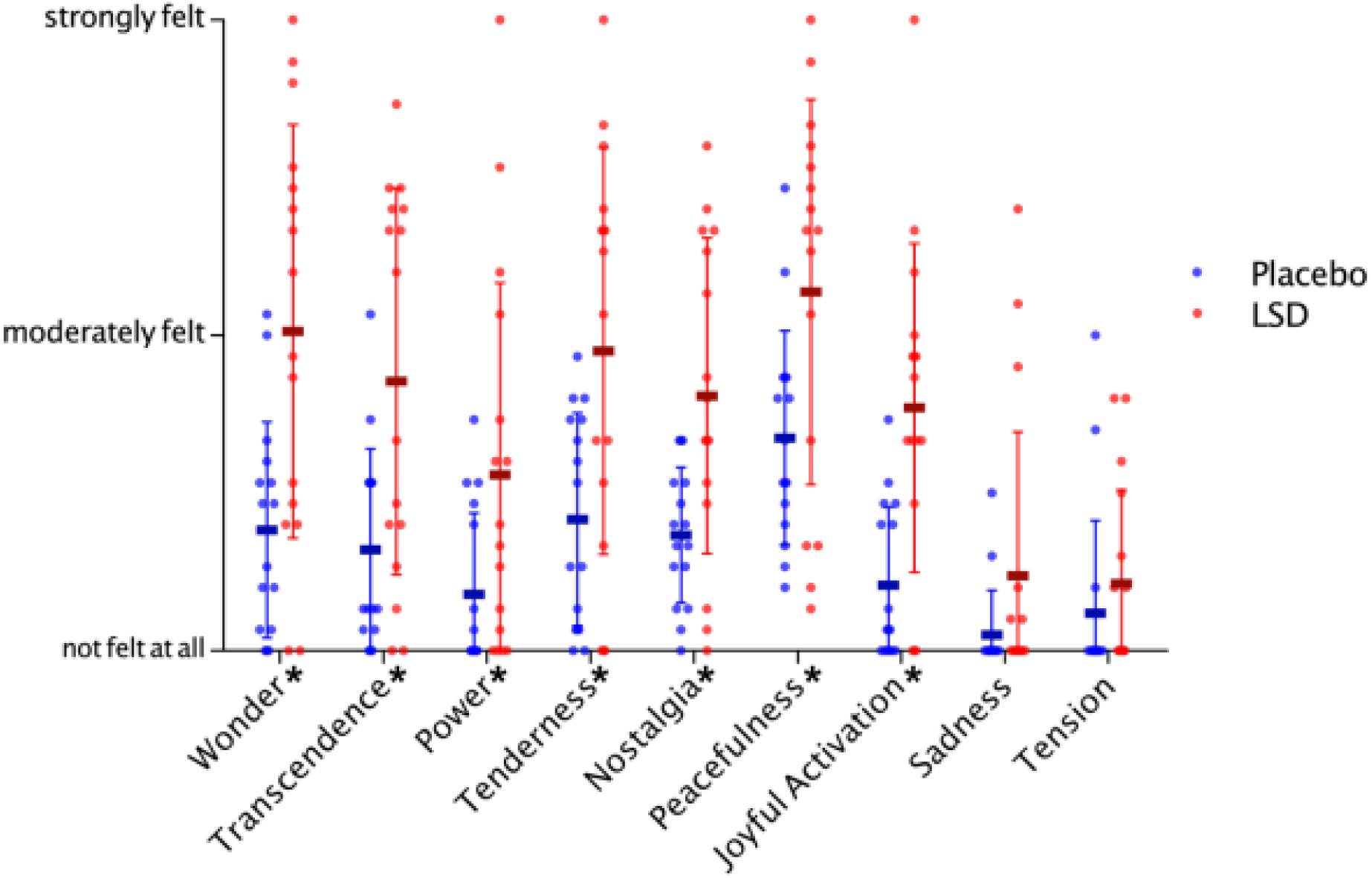
The average scores (thick line) + standard deviation (error bars) for each condition, including individual values (dots) for each participant, for nine factors of music-evoked emotions assessed with the 25-item Geneva Emotional Music Scale (GEMS). Blue = placebo and red = LSD. Asterisks on the x-axis by the factor labels indicate a statistically significant difference for this emotion between condition (p ≤ 0.05) after multiple comparison correction.

### 3.4 Correlation analyses between music-evoked brain activity and peak emotions

Having established that psychedelics intensify the emotional experience of music and alter the music-evoked BOLD response, we subsequently explored interactions between peak emotions (as captured by the GEMS scales *wonder* and *transcendence*) and associated increases in music-evoked brain activity. We conducted correlation analyses (Spearman’s rank) between changes in music-evoked peak emotions (LSD > Placebo) and music-evoked BOLD activity (LSD > Placebo) for nine ROIs. This correlation analysis was constrained to the contrast for timbral complexity (i.e., PC3) as related BOLD changes were most pronounced and occurred in brain networks commonly identified for music perception and emotion (Alluri et al., 2012; Koelsch, 2014). ROI selection was based on previous literature and encompassed structures that have been commonly identified for music-evoked emotion (striatum, precuneus and insula)(Koelsch, 2014; Trost et al., 2012), auditory perception (planum temporale and IFG) (Alluri et al., 2012, 2013; Griffiths and Warren, 2002; Zatorre and Salimpoor, 2013) and regions previously associated with the subjective effects of psychedelics (precuneus and insula) (Carhart-Harris et al., 2016a; Muthukumaraswamy et al., 2013a; Tagliazucchi et al., 2016). We found significant positive correlations between changes in music-evoked feelings of *wonder* and music-evoked BOLD activation to timbral complexity within the precuneus (*r* = 0.67, *p* = .034) as well as within the right inferior frontal gyrus (*r* = 0.65, *p* = .045) (Fig. 4). In addition, we found trend-level positive correlations between changes in feelings of *transcendence* and the right inferior frontal gyrus (*r* = 0.61, *p* = .066). P-values reported were corrected for multiple comparisons using one-sided permutation testing with α set to .05.

**Figure 4.**
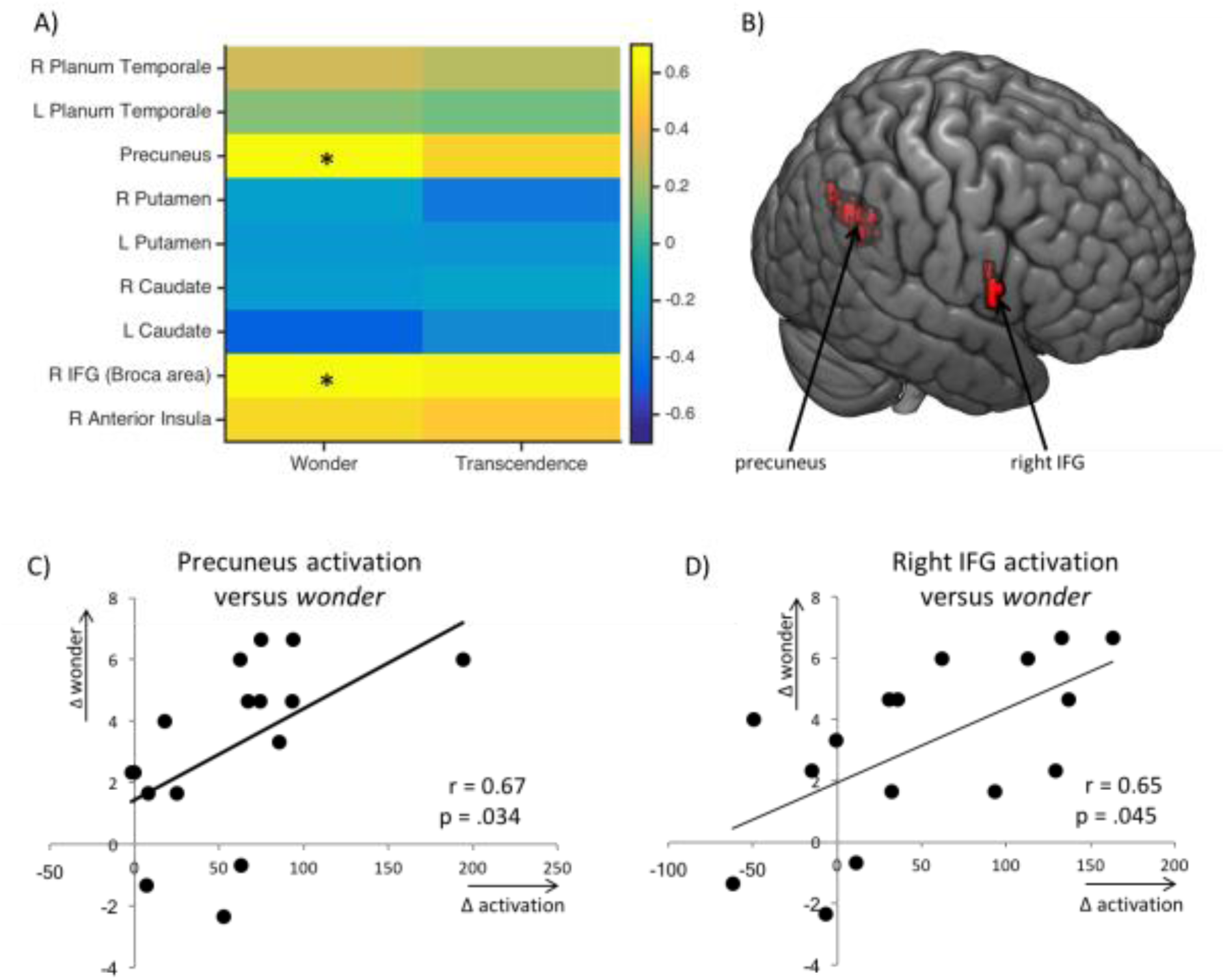
The correlation analysis comparing subjective experience with BOLD activity. **A)** Correlation matrix of parameter estimates with music-evoked peak emotions: wonder and transcendence. Asterisks indicate a statistically significant correlation (p 5 0.05) after multiple comparison correction. **B)** Location of masks for the precuneus and right inferior frontal gyrus (right IFG). **C)** Correlation plot showing a positive correlation between changes in parameter estimates in the precuneus and increases in music-evoked emotions of wonder under LSD (r = 0.67, p = .034). **D)** Correlation plot showing a positive correlation between changes in parameter estimates in the right IFG and increases in music-evoked emotions of wonder under LSD (r = 0.65, p = .045).

**Table 2.** MNI coordinates for the centres of gravity of ROIs

| Region | Coordinates (X,Y Z) |  |  | # voxels |
| --- | --- | --- | --- | --- |
| R Planum Temporale | 54.8 | -22.6 | 11 | 290 |
| L Planum Temporale | -48.8 | -33 | 15.2 | 658 |
| Precuneus | 14.6 | -62.6 | 19 | 414 |
| R Putamen | 22.8 | 10.2 | -0.4 | 308 |
| L Putamen | -25.6 | 1.4 | 3.8 | 364 |
| Right Caudate | 15.2 | 14 | 8 | 158 |
| Left Caudate | -14.2 | 17.8 | 3.4 | 76 |
| Right IFG (Broca area) | 58.8 | 9.8 | 14.6 | 60 |
| Right Anterior Insula | 31.2 | 13.8 | 9 | 48 |

### 3.5 Effects of psychedelics and music on functional connectivity

The relationship between feelings of wonder and music-evoked activation to timbral complexity in the precuneus and the right IFG informed subsequent PPI analyses with the aim of gaining insight into underlying music-specific changes in functional connectivity under LSD. Two separate PPI analyses were conducted, using either the precuneus or right IFG as a seed ROI, in order to identify brain areas for which the activation time course during the LSD condition was more or less coupled with the seed regions depending on the level of timbral complexity in the music. Under LSD, we found significantly increased coupling of the precuneus with the right superior frontal gyrus (SFG) for passages when the music exhibited high timbral complexity (See fig. 5A). In addition, we found significantly decreased coupling of the precuneus with the right IFG as well as the right auditory cortex under LSD when timbral complexity in the music increased (See fig. 5B). No significant timbral-specific changes in functional connectivity were found for the right IFG as seed.

**Figure 5.**
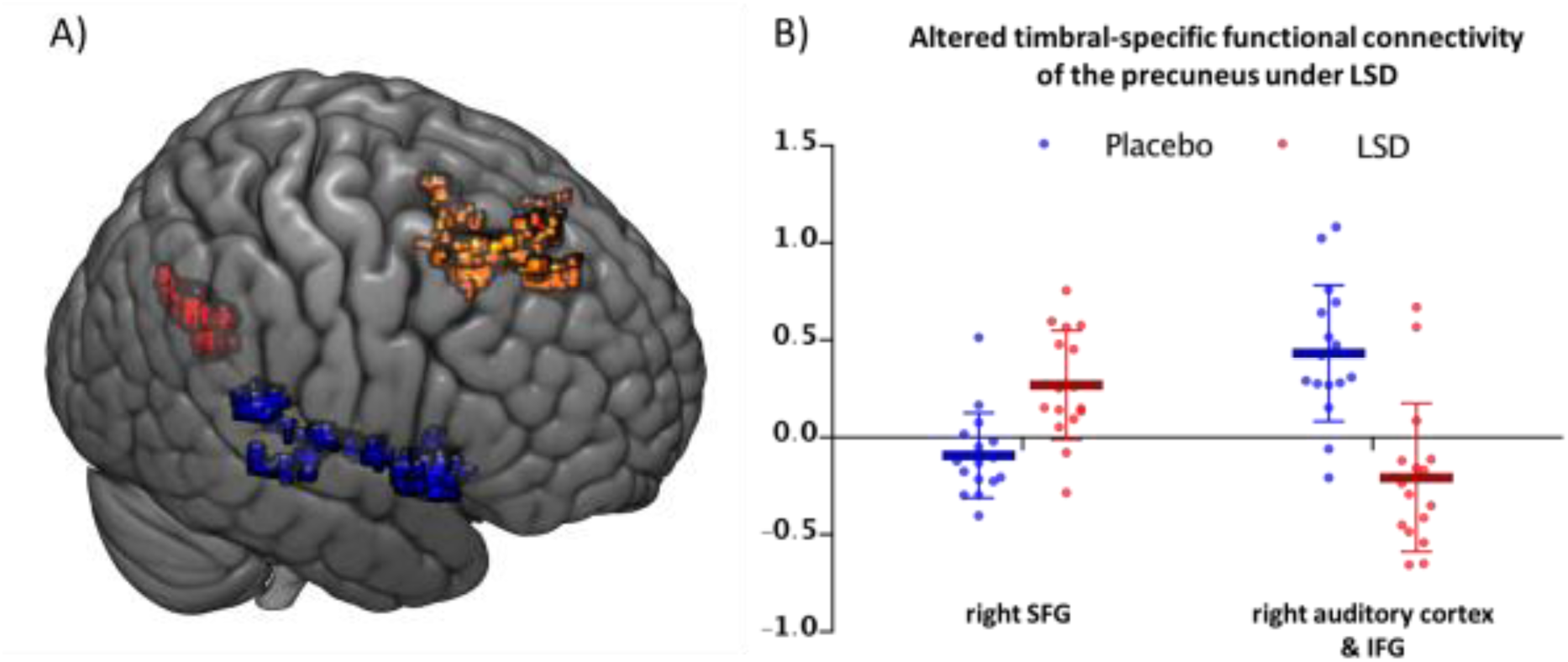
Brain regions showing altered timbral-specific functional connectivity under LSD. **A).** Group level results from the PPI analysis using the right precuneus ROI (displayed in red). Increased coupling (LSD>placebo) with the precuneus was observed in the right superior frontal gyrus (right SFG) (displayed in hot/orange), and decreased coupling (placebo>LSD) with the precuneus was observed in the right auditory cortex and right inferior frontal gyrus (right IFG) (displayed in blue). Z < 2.3. **B)** Connectivity values of each individual participant and for each condition (Blue = placebo, red = LSD) for the precuneus with the right SFG (left), and for the precuneus with the right IFG and auditory cortex (right).

### 3.6 Effect of head motion

In order to investigate potentially confounding effects of motion on our results, we investigated frame-wise displacement (FD) between the drug and placebo condition as well as conducted correlation analyses between changes in FD (LSD>placebo) and extracted parameter estimates from the nine ROIs. The mean ± std FD for placebo and LSD was 0.091 ± 0.037 and 0.174 ± 0.1, respectively, which was found to be statistically significant using a paired t-test (*t*_(15)_ = −3.28, *p*=0.005). However, we did not find any significant correlations (Spearman’s rank) between changes in FD (LSD > Placebo) and changes in music-evoked BOLD activity for the contrast timbral complexity (LSD > Placebo) in any of the nine ROIs: right planum temporale (*r*=0.04, *p*=0.88), left planum temporale (*r*=-0.01, p=0.97), precuneus (*r*=0.02, *p*=0.93), right putamen (*r*=−0.29, *p*=0.27), left putamen (*r*=-0.21, *p*=0.42), right caudate (*r*=-0.2, *p*=0.45), left caudate (*r*=-0.28, *p*=0.3), right IFG (*r*=0.29, *p*=0.27) and right insula (*r*=0.28, *p*=0.28). We therefore concluded that the effect of motion for our results seem negligible. P-values reported here are uncorrected for multiple comparisons.

## 4. Discussion

This study investigated the effects of the classic psychedelic drug LSD on music-evoked brain activity and emotion under naturalistic listening conditions. Widespread BOLD activity changes under LSD in cortical and subcortical areas were associated with distinct acoustic features in the music. The BOLD signal increases observed under LSD to the component timbral complexity are particularly noteworthy, as the regions lie within brain networks commonly identified for music-perception and music-evoked emotion. These include bilateral auditory cortices, right inferior frontal gyrus (IFG), right insula, precuneus, bilateral striatum and supplementary motor area (SMA). Music-evoked emotions were rated higher under LSD, and increased feelings of wonder correlated with increased BOLD activation to timbral complexity in the right precuneus and in the right IFG. The functional connectivity analysis revealed decoupling under LSD of the right precuneus with a network that includes auditory cortices and the right IFG with increased timbral complexity, as well as increased coupling under LSD of the precuneus with the superior frontal gyrus (SFG).

### 4.1 Timbral complexity and the auditory-IFG network

Timbral complexity corresponds to the complexity in the shape and the spread of the music’s spectral envelope. Spectral properties are thought to be processed by the planum temporale (Kumar et al., 2007), a central “computational hub” of the auditory cortex that segregates incoming spectral signals into distinct spectrotemporal patterns (Griffiths and Warren, 2002). Subsequent information exchange with higher cortical regions such as the IFG, is thought to facilitate the linking of the perceived spectrotemporal patterns with learned representations and associations (Hackett, 2011; Zatorre and Salimpoor, 2013). The auditory cortex and IFG are anatomically (Plakke and Romanski, 2014) and functionally (Tomasi and Volkow, 2012) well-connected, and considered an important pathway for the perception of auditory objects (Griffiths and Warren, 2004; Zatorre and Salimpoor, 2013), musical melody (Zatorre et al., 1994), musical harmony (Opitz et al., 2002; Schönwiesner et al., 2007), musical syntax (Koelsch et al., 2002), and the encoding of musical expectation (Osnes et al., 2012; Zatorre, 2015) as well as musical imagination (Herholz et al., 2012). Damage and abnormal functioning in the auditory-IFG network has been linked to impaired music perception (Hyde et al., 2007, 2011; Leveque et al., 2016) and auditory hallucinations (Looijestijn et al., 2013). Therefore, the marked BOLD signal changes under LSD to timbral complexity within this network suggests how LSD may modulate the brain’s processing of timbral and spectrotemporal information in music.

Apart from a key role in music perception, converging evidence describes the IFG’s involvement in the evaluation of emotional valence in acoustic information. Happy emotional vocalisations produce increased IFG activity compared to angry vocalisations (Frühholz et al., 2016; Schirmer and Kotz, 2006; Wiethoff et al., 2008) - an effect that has been observed as early in development as 7-months old (Grossmann et al., 2010). The brain has been shown to more closely track the time-varying tonal structure of music in the IFG during the experience of music-evoked nostalgia than during the experience of music that does not evoke nostalgia (Barrett and Janata, 2016). Brain damage and experimentally disrupting brain activity in the IFG, by using transcranial magnetic stimulation (TMS), is linked with impairments in identifying emotional valence in voice (Hoekert et al., 2010). However, the IFG is associated with emotionally salient information from different sensory modalities (Lee and Siegle, 2012), and has been more generally conceptualised as being part of a cognitive control network (Irlbacher et al., 2014; Marklund and Persson, 2012) that is associated with the allocation of attentional resources for emotionally salient information (Vuilleumier, 2005). Paying attention to the emotional valence of visual (Critchley et al., 2000; Schindler and Kissler, 2016) and auditory (Bach et al., 2008) stimuli is associated with increases in IFG activity, and emotionally salient information can produce marked attentional biases (Vuilleumier, 2005). Increased right IFG activation to music’s timbral complexity under LSD, and the positive correlation of this increase with music-evoked emotion, may therefore indicate increased allocation of attentional resources for emotional information conveyed via music’s timbre that is processed within the auditory-IFG circuitry.

The view that the auditory-IFG network may play a role in LSD’s effects on musical information processing receives further support from the functional connectivity analyses. The PPI analysis revealed significant decoupling of the right precuneus with the right auditory cortex and the right IFG under LSD, in interaction with timbral complexity. The precuneus is a highly connected hub region that plays an important role in self-referential cognition (Buckner et al., 2008; Cavanna and Trimble, 2006) and emotion-regulation (Cavanna and Trimble, 2006; Schilbach et al., 2012), and shows desynchronized activity under psychedelics (Carhart-Harris et al., 2016a; Muthukumaraswamy et al., 2013a; Riba et al., 2002). This disorganising effect of psychedelics on precuneus activity has been associated with “ego-dissolution”, characterised by increased emotional lability (Carhart-Harris et al., 2016a; Lebedev et al., 2015; Tagliazucchi et al., 2014, 2016). Although the precuneus is activated by music-evoked emotion (Baumgartner et al., 2006; Koelsch, 2014; Trost et al., 2012), the region shows poor anatomical (Cavanna and Trimble, 2006) and resting state functional connectivity (Tomasi and Volkow, 2012) with auditory regions. The precuneus is, however, well-connected with frontal regions such as the IFG and SFG (Zhang and Li, 2012), and this circuit is thought to provide top-down guidance (Cavanna and Trimble, 2006). As such, decreased precuneus-auditory/IFG coupling under LSD may reflect a reduced regulatory influence of the precuneus over emotion processing within the IFG, and this in turn may facilitate an intensified emotional response to music.

### 4.2 Timbral complexity and the limbic system

LSD produced marked increases in BOLD activation to timbral complexity in the striatum and in the insula; regions that have both been consistently associated with music-evoked emotion. The striatum has been implicated in brain mechanisms underlying music-evoked pleasure (Blood and Zatorre, 2001; Salimpoor et al., 2011), likely via the encoding of auditory expectation and reward in interaction with auditory cortices (Salimpoor et al., 2013, 2015). The insula is a high-level hub region that is thought to facilitate emotional awareness (Craig, 2009), the attribution of emotional meaning to music, sound and human voice (Frühholz et al., 2016; Koelsch, 2014; McGettigan et al., 2013; Petrini et al., 2011; Trost et al., 2012), and the subjective effects of classic psychedelics(Carhart-Harris et al., 2016a; Lebedev et al., 2015; Tagliazucchi et al., 2014, 2016). Interestingly, the present study did not find a relationship between music-evoked emotion and the striatum. One implication of this could be that the striatum is predominantly concerned with the processing of “simple” affect states characterised by reward and pleasure, whereas more complex emotional states, such as feelings of wonder, may require more complex cognition, and thus more complex cortical mechanisms. The selective positive correlation between increased feelings of wonder and BOLD signal changes in higher-level brain regions such the IFG and precuneus support this idea.

Taken together, these observations suggest that LSD alters the dynamic interplay between the analysis of “low-level” spectral properties in the music, and the subsequent “high-level” regulation and processing of the emotional associations. The precuneus, the IFG and the insula may play a particularly important role in facilitating music-evoked peak emotion, due to their involvement in processing of emotional awareness (Craig, 2009) and “complex” music-evoked emotion (Frühholz et al., 2016; Koelsch, 2014; McGettigan et al., 2013; Petrini et al., 2011; Trost et al., 2012). A disintegration or weakening of the mechanisms that normally regulate emotion (Lebedev et al., 2015; Tagliazucchi et al., 2016), may be one mechanism underlying the strong emotional experiences associated with music under psychedelics.

### 4.3 Possible brain mechanisms

Classic psychedelics such as LSD stimulate serotonin 2A receptors (Kometer et al., 2013; Riga et al., 2016; Titeler et al., 1988), which are primarily expressed on deep layer V pyramidal cells (Celada et al., 2013). Upon activation of serotonin 2A receptors, the cell’s membranes depolarize, reducing the neuron’s threshold for firing (Andrade, 2011). Over-activation of the serotonin 2A receptor underlies desynchronised activity within high-level brain networks (Muthukumaraswamy et al., 2013b), leading to impaired top-down control and an increased information exchange between brain modules that are usually more strictly functionally segregated (Carhart-Harris et al., 2016a; Tagliazucchi et al., 2016). Deep layer V pyramidal cells are believed to encode learned mental representations or “predictions” that are matched with incoming sensory stimuli in a top-down fashion (Bastos et al., 2012), and a disruption of this process via over-activation of the serotonin 2A receptor system by psychedelics is believed to produce disorganised or “entropic” activation of these mental representations (Carhart-Harris et al., 2014).

The high density of serotonin 2A receptors within the precuneus, insula and the planum temporale (Erritzoe et al., 2009; Ettrup et al., 2014) suggests that psychedelics can alter auditory information processing at different stages of the processing hierarchy. Since the planum temporale is functionally specialised for processing distinct spectrotemporal patterns (Griffiths and Warren, 2002), the particularly pronounced BOLD increases under LSD for “timbral complexity” in this region may reflect an increased activation of intrinsic deep-pyramidal neurons that encode predictions for music’s spectral properties. In addition, altered planum temporale activity may also be driven by increased input from top-down projecting deep layer V pyramidal cells from the IFG and insula. In turn, decoupling of the auditory/IFG circuitry with the precuneus could indicate an unregulated (“entropic”) activation of emotional associations attributed to the music by this circuit (Frühholz et al., 2016; Koelsch, 2014; McGettigan et al., 2013; Petrini et al., 2011; Trost et al., 2012), hence yielding an intensified emotional response to music.

### 4.4 The significance of timbre

This discussion leads to the question of why LSD produced especially prominent modulation of neural processing of music’s timbral complexity. Timbre-perception is present in early infancy (Dalla Bella et al., 2001), and the infants’ ability to memorise and recognize their mother’s voice as early as two days after birth has been argued to be due to the infant’s innate tuning to timbral properties common to speech and music (Tsang and Trainor, 2002). Timbre is shown to convey emotion cross-culturally (Egermann et al., 2015), independently of other acoustic features (Hailstone et al., 2009). Timbre has been conceptualised to function as an interface, that connects distinctive spectral features within a sound with meaningful emotional associations (Hailstone et al., 2009). The findings of this study suggest that LSD may target this interface, thereby “tuning” the brain to acoustic properties that communicate emotionally significant information. The resulting heightened responsiveness for music’s “non-verbal language” may be at the core of the intensifying effects of LSD for music-evoked emotion.

### 4.5 Effects of LSD on the brain’s processing of other acoustic features

The previous discussion centred around the changes observed for timbral-complexity, because these changes were most pronounced and appeared to be localised to networks commonly associated with music-perception and music-evoked emotion. However, several other findings must be highlighted. Increased thalamus and decreased IFG activation was observed under LSD in response to “fullness”, i.e., the loudness of the sound. Loudness reflects a less complex acoustic property (the amplitude of the signal), and has previously been associated with a preferential activation of lower auditory regions (Reiterer et al., 2007). Significant BOLD signal changes were also observed in occipital cortices for various acoustic features. Since the serotonin 2A receptor is densely expressed in the visual cortex and LSD produces significant alterations in the intrinsic functional organisation of the visual cortex (Roseman et al., 2016), these occipital changes could reflect effects of LSD on music-induced mental imagery, such as reported elsewhere (Kaelen et al., 2016).

### 4.6 Therapeutic implications

Music listening is considered an important component of psychedelic-assisted psychotherapy for the treatment of depression (Carhart-Harris et al., 2016b), addiction (Bogenschutz et al., 2015; Johnson et al., 2014) and end-of-life care (Gasser et al., 2014; Griffiths et al., 2016; Grob CS et al., 2011; Ross et al., 2016). The present study supports the usefulness of music in facilitating emotions associated with peak-experiences, which are associated with positive therapy outcomes (Garcia-Romeu et al., 2014; Griffiths et al., 2016; Ross et al., 2016), and is the first to ground this framework into a neurobiological basis. The findings also suggest that knowledge of the ways musicians utilise timbre to convey emotion may be a particularly fruitful avenue to empirically inform the design of playlists for psychedelic-assisted psychotherapy.

### 4.7 Study limitations

The study focused on LSD’s effects on the neurophysiological processing of acoustic properties in music, but did not include perceptual measures for these features. Future studies may address questions such as whether features are perceived differently, whether certain acoustic features are experienced differently than others and whether perceptual changes reflect real acoustic details that have been previously unnoticed. Studies assessing these issues will be particularly important for our understanding of how psychedelics alter the subjective experience of music, and more broadly, salient sensory information.

Another study limitation was that only a single musical genre was used. The music used was rich in timbre and pitch variation, but low in tempo. The music choice may therefore have resulted in restricted description of LSD’s effect on timbral features in music, since it could be hypothesised that other brain networks will be engaged when more rhythmic and arousing music is used. However, the component for which LSD showed most pronounced neurophysiological changes (i.e. “timbral complexity”), was not the component that explained most of the variance within the music (i.e., the third component), supporting the validity of the method in identifying brain changes to distinct and diverse musical dynamics within a naturalist music listening experience.

### 5. Conclusion

The present study revealed altered brain activity and connectivity to acoustic features in music under LSD. Most pronounced changes in brain activity and connectivity were associated with the component timbral complexity, representing the complexity of the music’s spectral distribution, and these occurred in brain networks previously identified for music-perception and music-evoked emotion, and showed an association with enhanced music-evoked feelings of wonder under LSD. The findings are the first to shed light on how the brain processes music under LSD, and provide a new perspective on brain processes underlying music perception and peak emotion under naturalistic listening conditions.

## Acknowledgements

This research received financial and intellectual support from the Beckley Foundation (Grant number: P41825) and was conducted as part of a wider Beckley-Imperial research programme. The researchers would like to thank supporters of the Walacea.com crowd-funding campaign who played a crucial role in securing funds to complete the study.

## Footnotes

1 The term peak experience was firstly coined by the psychologist Abraham Maslow (Maslow, 1964, 1971). Peak-experiences are characterized by a transcending of the usual sense of self, and are typically accompanied by feelings of wonder, bliss, union and awe. Peak experiences have also been referred to as *mystical-type* of *spiritual* experiences, and can occur under natural conditions such as when being deeply moved by a work of art or a moment of creative inspiration but are frequently reported with psychedelics (Garcia-Romeu et al., 2014; Griffiths et al., 2016; Ross et al., 2016)

